# Immune escape of SARS-CoV-2 Omicron variant from mRNA vaccination-elicited RBD-specific memory B cells

**DOI:** 10.1101/2021.12.21.473528

**Authors:** Aurélien Sokal, Matteo Broketa, Annalisa Meola, Giovanna Barba-Spaeth, Ignacio Fernández, Slim Fourati, Imane Azzaoui, Andrea de La Selle, Alexis Vandenberghe, Anais Roeser, Magali Bouvier-Alias, Etienne Crickx, Laetitia Languille, Marc Michel, Bertrand Godeau, Sébastien Gallien, Giovanna Melica, Yann Nguyen, Virginie Zarrouk, Florence Canoui-Poitrine, France Noizat-Pirenne, Jérôme Megret, Jean-Michel Pawlotsky, Simon Fillatreau, Etienne Simon-Lorière, Jean-Claude Weill, Claude-Agnès Reynaud, Félix A. Rey, Pierre Bruhns, Pascal Chappert, Matthieu Mahévas

**Affiliations:** Institut Necker Enfants Malades (INEM), INSERM U1151/CNRS UMR 8253, Université de Paris, Paris, France; Institut Pasteur, Université de Paris, INSERM UMR1222, Unit of Antibodies in Therapy and Pathology, Paris, France; Institut Pasteur, Université de Paris, CNRS UMR 3569, Unité de Virologie Structurale, Paris, France; Département de Virologie, Bactériologie, Hygiène et Mycologie-Parasitologie, Centre Hospitalier Universitaire Henri-Mondor, Assistance Publique-Hôpitaux de Paris (AP-HP), Créteil, France; INSERM U955, équipe 18. Institut Mondor de Recherche Biomédicale (IMRB), Université Paris-Est Créteil (UPEC), Créteil, France; Service de Médecine Interne, Centre Hospitalier Universitaire Henri-Mondor, Assistance Publique-Hôpitaux de Paris (AP-HP), Université Paris-Est Créteil (UPEC), Créteil, France; INSERM U955, équipe 2. Institut Mondor de Recherche Biomédicale (IMRB), Université Paris-Est Créteil (UPEC), Créteil, France; Service de Maladies Infectieuses, Centre Hospitalier Universitaire Henri-Mondor, Assistance Publique-Hôpitaux de Paris (AP-HP), Université Paris-Est Créteil (UPEC), Créteil, France; Service de Médecine interne, Hôpital Beaujon, Assistance Publique-Hôpitaux de Paris, Université de Paris, Clichy, France; Département de Santé Publique, Unité de Recherche Clinique (URC), CEpiA (Clinical Epidemiology and Ageing), EA 7376- Institut Mondor de Recherche Biomédicale (IMRB), Centre Hospitalier Universitaire Henri-Mondor, Assistance Publique-Hôpitaux de Paris (AP-HP), Université Paris-Est Créteil (UPEC), Créteil, France; Etablissement Français du Sang (EFS) Ile de France, Créteil, France; Plateforme de Cytométrie en Flux, Structure Fédérative de Recherche Necker, INSERM US24-CNRS UMS3633, Paris, France; Institut Pasteur, Université de Paris, Evolutionary genomics of RNA viruses, Paris, France

**Author notes:** these authors contributed equally (Aurélien Sokal, Matteo Broketa, Annalisa Meola). these authors contributed equally (Giovanna Barba-Spaeth, Ignacio Fernández, Slim Fourati, Imane Azzaoui, Andrea de La Selle, Alexis Vandenberghe). shared senior authorship (Jean-Claude Weill, Claude-Agnès Reynaud, Félix A. Rey, Pierre Bruhns, Pascal Chappert, Matthieu Mahévas). Lead Contact: Matthieu Mahévas.

## Abstract

Memory B cells (MBCs) represent a second layer of immune protection against SARS-CoV-2. Whether MBCs elicited by mRNA vaccines can recognize the Omicron variant is of major concern. We used bio-layer interferometry to assess the affinity against the receptor-binding-domain (RBD) of Omicron spike of 313 naturally expressed monoclonal IgG that were previously tested for affinity and neutralization against VOC prior to Omicron. We report here that Omicron evades recognition from a larger fraction of these antibodies than any of the previous VOCs. Additionally, whereas 30% of these antibodies retained high affinity against Omicron-RBD, our analysis suggest that Omicron specifically evades antibodies displaying potent neutralizing activity against the D614G and Beta variant viruses. Further studies are warranted to understand the consequences of a lower memory B cell potency on the overall protection associated with current vaccines.

## Introduction

SARS-CoV-2 Omicron has only recently emerged but is already starting to overcome the previously dominant Delta lineage in many countries, suggesting a strong selective advantage. The Spike protein of SARS-CoV-2 Omicron harbors 32 mutations as compared to the original Wuhan strain, with particular hotspots of mutations in the ACE2 receptor binding domain (RBD) (15 amino acid substitutions) and in the N-terminal domain (NTD) (3 deletions, 1 insertion, 4 substitutions). Of particular concern, the Omicron variant displays key mutations previously associated with immune escape (K417N, E484A, T478K in the RBD) or enhanced infectivity (N501Y, P681H), but also numerous mutations rarely detected in previous variants. SARS-CoV-2 Omicron may thus have emerged after extensive selection based on beneficial combinatorial effects, as was predicted in silico for the Q498R mutation for example (Zahradník et al., 2021). The overall mutational profile of the Omicron variant thus suggests both increased immune escape and increased infectivity.

Despite sizeable immune evasion by some of the previous variants of concern (VOCs), mRNA vaccines have so far maintained strong protection in recently vaccinated individuals due to an initially broad and strong serum IgG response. This response is nonetheless waning with time. We and others have been able to show that SARS-CoV-2-specific memory B cells (MBCs) represents a potent layer of additional immune protection (Dugan et al., 2021; Gaebler et al., 2021; Rodda et al., 2021; Sokal et al., 2021a, 2021b). MBCs not only persist after infection but continuously evolve and mature by progressive acquisition of somatic mutations in their variable region genes to improve affinity through an ongoing germinal center response (Gaebler et al., 2021, 2021; Rodda et al., 2021; Sokal et al., 2021a). Upon restimulation, either in the context of natural infection or vaccinal boost, MBCs can rapidly differentiate into the plasma cell lineage, secreting the diverse array of high-affinity antibodies contained in the their repertoire. Deep analysis of the repertoire of vaccinated individuals has so far suggested that a sizeable proportion of such MBCs is able to neutralize all VOCs up to the Beta variant (Sokal et al., 2021b; Wang et al., 2021). A non-peer reviewed preprint suggests that SARS-CoV-2 Omicron escapes to a large extent vaccine-elicited antibodies or antibodies from SARS-CoV-2 recovered sera (Planas et al., 2021). Yet, the intrinsic capacity of the MBC pool to recognize Omicron variant still remains unexplored.

## Results

We recently performed in depth characterization of over 400 single-cell sorted and cultured RBD-specific MBCs isolated from 8 vaccinated SARS-CoV-2 recovered and 3 vaccinated naive individuals (Sokal et al., 2021b). Our analyses included affinity measurements against VOCs and VOIs (B.1.1.7, Alpha; B.1.351, Beta; P.1, Gamma; B.1.617.2, Delta; B.1.617.1 Kappa) and neutralization potency against the D614G and Beta SARS-CoV-2 variants. Selected individuals were part of two longitudinal cohorts, one of COVID-19 patients (MEMO-COV2 (Sokal et al., 2021a)) followed for one year after initial infection prior to vaccination, and one of healthcare workers, with no clinical history of COVID-19 and no serological evidence of previous SARS-CoV-2 infection, vaccinated as part of the French vaccination program. All individuals had received the BNT162b2 vaccine and were sampled for circulating MBC analyses 7 days and 2 months after boost (Sokal et al., 2021b). In all individuals, we were able to detect a sizeable fraction of MBCs encoding antibodies with high affinity and neutralizing potential against all tested VOCs, with the Beta variant showing the largest extent of immune escape at the time. This suggested that MBC elicited by prior infection or vaccination would be able to provide an efficient secondary layer of protection in case of waning of the serological protective antibodies or escape of a novel SARS-CoV-2 variant from this antibody pool.

To test whether such conclusion still holds true in the context of the newly emerging B.1.1.529 - Omicron variant, we generated a recombinant Omicron RBD harboring the 15 described mutations, including N501Y, K417T/N, E484K/Q/A, T478K found in other VOCs and additional mutations G339D, S371L, S373P, S375F, N440K, G446S, S477N, Q493R, G496S, Q498R, Y505H (**Figure 1A, B**). All previously described naturally expressed monoclonal IgG from single-cell culture supernatants of MBCs were assayed anew using Biolayer interferometry (BLI) to assess their affinity against the Omicron RBD, and 313 monoclonal supernatants passed quality controls. As previously reported, monoclonal antibodies encoded by MBCs isolated from vaccinated COVID-19-recovered patients and vaccinated naive patients contained a vast majority of high-affinity binders against the ancestral Wuhan (WT) RBD, among which ~50% remained of high-affinity against Beta and Delta RBD. This proportion further decreased against Omicron RBD to ~30% (**Figure 1C**). Importantly, ~40% of the clones showed no or low-affinity (KD>10^-8^ M) to the Omicron RBD, in both vaccinated recovered and vaccinated naive individuals, in line with the prediction that Omicron accumulates numerous mutations associated with antibody evasion. Similar conclusions could be drawn when focusing only on the highly mutated antibodies found in recovered individuals (**Figure 1D-E**). This further suggests that the affinity maturation process, which takes place in the germinal centers of both infected and vaccinated individuals and selects over time for high-affinity clones (KD<10^-9^ M) against the Wuhan (WT) RBD (**Figure 1D**), is of lower benefit in the context of such a highly mutated variant like Omicron.

**Figure 1.**
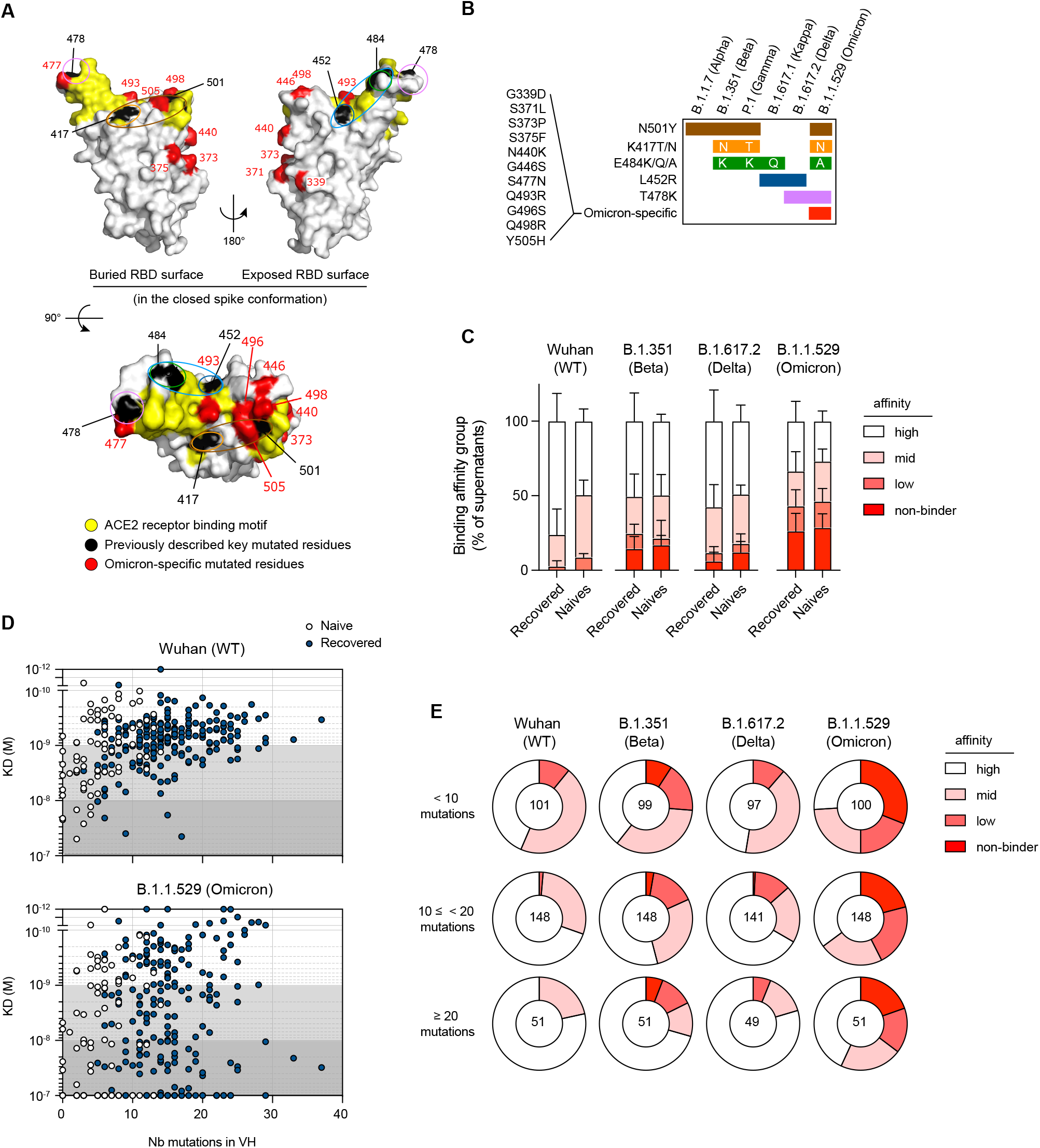
The memory B cell pool of vaccinated individuals contains a reduced frequency of high-affinity clones against the Omicron RBD. **(A)** RBD (extracted from the PDB:6XR8 spike protein trimer structure) shown in three orthogonal views with the ACE2 receptor binding motif highlighted in yellow and the residues found mutated in at least one of the Alpha, Beta, Gamma, Kappa or Delta variants - L452, K417, T478, E484 and N501-highlighted in black. Residues specifically mutated in omicron - G339, S371, S373, N440K, G446, S477, Q493, G496, Q498, Y505H are highlighted in red. Single or group of mutations predicted as key binding residues for particular antibodies are further highlighted by colored ovals according to the color scheme used in (B) and Figure 2B. **(B)** Distribution of known mutations in the RBD domain between B.1.1.7 (Alpha), B.1.351 (Beta), P.1 (Gamma), B.1.617.1 (Kappa), B.1.617.2 (Delta) and B.1.1.529 (Omicron) SARS-CoV-2 variants. **(C)** Histogram showing the binding affinity distribution of monoclonal antibodies from single-cell culture supernatants of RBD-specific MBCs isolated from vaccinated SARS-CoV-2 recovered (n=225) and vaccinated naive donors (n=88) against the Wuhan (WT) RBD, B.1.351 (Beta), B.1.617.2 (Delta) and B.1.1.529 (Omicron) RBD variants, defined as: high (KD <10^-9^ M), mid (10^-9^ ≤ KD <10^-8^ M) and low (10^-8^≤ KD <10^-7^). Bars indicate mean±SEM. **(D)** Measured KD (M) against Wuhan (WT) (upper panel) or B.1.1.529 (Omicron) RBD (bottom panel) vs. number of V_H_ mutations for all tested monoclonal antibodies with available V_H_ sequence from SARS-CoV-2 recovered (dark blue) and naive (white) donors (Spearman correlations for all sequences: V_H_ mutation/WT KD: r = 0.3791, P<0.0001; V_H_ mutation/B.1.1.529 KD: r = 0.1597, P=0.0058). **(E)** Pie chart showing the binding affinity distribution of all tested monoclonal antibodies grouped according to their overall number of mutations: low (<10 mutations, upper panel), intermediate (<20 and ≥10, mid panel) or high V_H_ mutation numbers (≥20, lower panel). Affinity groups against Wuhan (WT), B.1.351 (Beta), B.1.617.2 (Delta) and B.1.1.529 (Omicron) RBD variants as defined in **(C)** are shown. Numbers at the center of each pie chart indicate the total number of tested monoclonal antibodies in each group.

Direct comparison of binding affinities towards the Wuhan (WT) and the Omicron RBD variant showed that ~50% (159/313) of monoclonal antibodies had reduced binding to Omicron RBD compared to Wuhan (WT) RBD (**Figure 2A**). Additional two by two comparisons of binding affinities towards the Wuhan (WT) and the previous RBD variants for these Omicron-affected antibodies further suggested that a large fraction of these antibodies is uniquely affected by the Omicron variant (**Figure 2A**). The distribution of mutations in the previous RBD variants (**Figure 1A and B**) had allowed us to predict the identity of key binding amino acid residues within the RBD for 125 MBC-derived monoclonal antibodies, affected by at least one of the previous RBD variants, out of the 313 included in this analysis (**Figure 2B** and (Sokal et al., 2021b)). 101 of these 125 MBC-derived monoclonal antibodies, including most of the ones recognizing the E484, K417 and N501 residues appeared also affected in their recognition of the Omicron variant. In addition, 58 out of the 188 previously unaffected antibodies appeared to be selectively affected by the new mutated residues uniquely displayed by Omicron (Omicron-specific, **Figure 2B**). Interestingly, clones recognizing the Omicron-specific residues were not enriched for the recurrent and convergent antibody rearrangements (IGHV1-2, IGHV1-69, IGHV3-53 and IGHV3-66; **Figure 2C**) observed in convalescent and vaccinated individuals (Barnes et al., 2020). Therefore Omicron-specific mutations appear to affect a more diverse range of VH recognizing the RBD than other VOCs and VOIs do.

**Figure 2.**
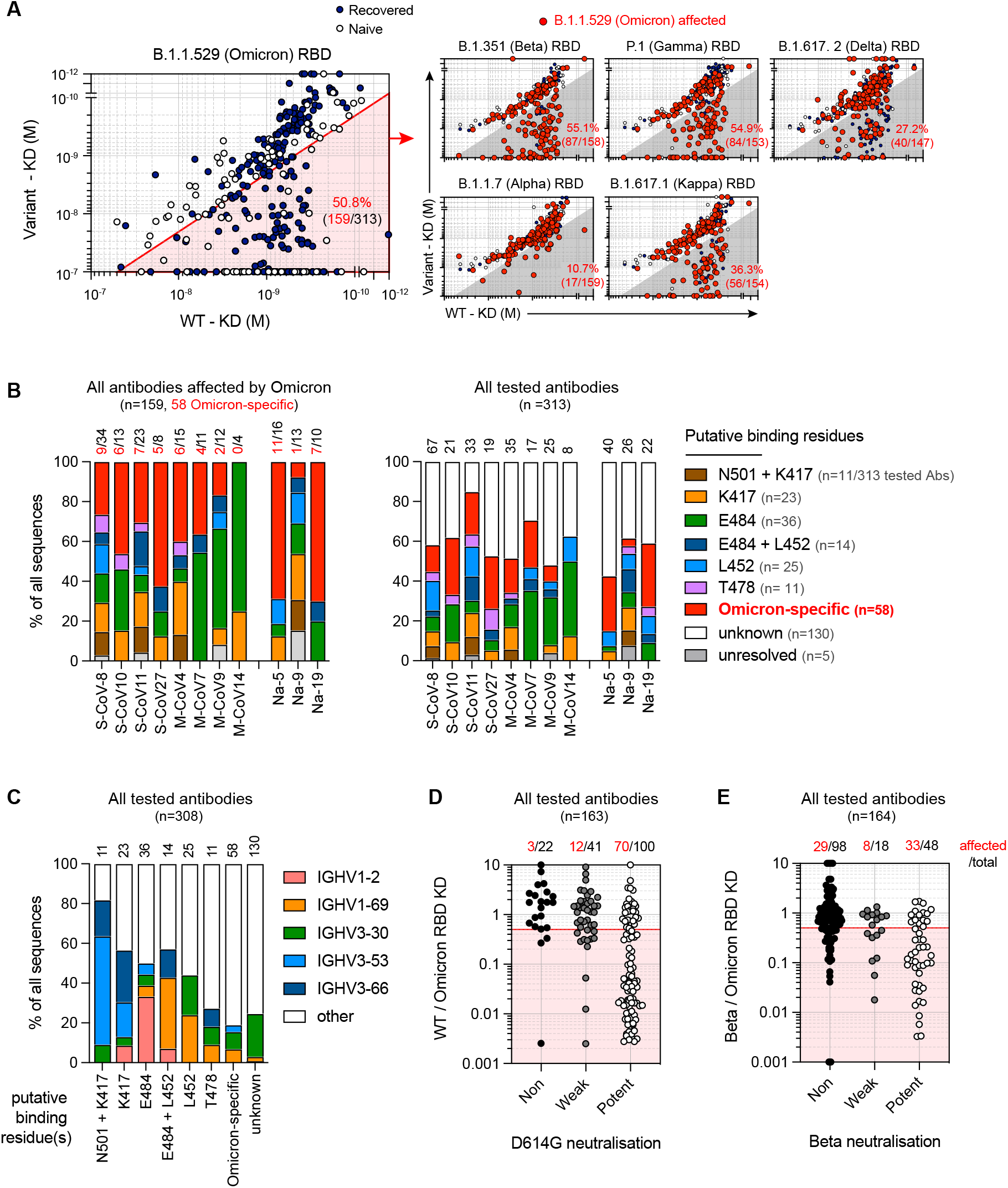
Omicron-specific RBD mutations expand its overall escape of memory B cell-derived antibodies. **(A)** Dot plot representing the KDs for B.1.1.529 (Omicron) RBD versus Wuhan (WT) RBD for all tested monoclonal antibodies from SARS-CoV-2 recovered (dark blue dots) and naive donors (white dots). The red shaded zone indicates monoclonal antibodies with at least two-fold increased KD for B.1.1.529 than for Wuhan (WT) (termed “B.1.1529-affected antibodies” herein) (left large panel). Dot plots representing the KDs for B.1.1.7 (Alpha), B.1.351 (Beta), P.1 (Gamma), B.1.617.1 (Kappa) and B.1.617.2 (Delta) RBD versus Wuhan (WT) RBD for all tested antibodies. B.1.1.529-affected antibodies are highlighted as larger size red dots (corresponding to clones present in the red sector in the left panel). Percentages indicate the proportion of B.1.1.529-affected monoclonal antibodies also affected by the indicated RBD variant. **(B)** Frequencies of antibodies targeting one of the predicted essential binding residue groups in Omicron-affected antibodies (left panel) and in all tested antibodies (right panel), as defined by RBD variants recognition profile in BLI, among all tested antibodies for each of the 11 individuals from whom memory B cells were assayed. Numbers of monoclonal antibodies for each donor are indicated on top of each histogram in black. Number of antibodies affected by Omicron specific mutations are detailed in red. **(C)** Proportion of IGHV1-2, IGHV1-69, IGHV3-30, IGHV3-53 and IGHV3-66 usage among all tested monoclonal antibodies with available V_H_ sequence and grouped based on their predicted essential binding residues, as defined in (B). Numbers of tested monoclonal antibodies for each donor are indicated on top of each histogram. (**D** and **E**) Ratio of Wuhan (WT) over B.1.1.529 RBD KD (D) or B.1.351 over B.1.1.529 KD for all monoclonal antibodies tested, grouped based on their neutralization potency against D614G **(D)** or B.1.351 SARS-CoV-2 (**E**) (Refer to Sokal et al. 2021b). Numbers on top indicate the numbers of monoclonal antibodies with a [KD for WT RBD / KD for B.1.1.529 RBD] ratio <0.5 (D) or a [KD for B.1.351 / KD for B.1.1.529 RBD] ratio <0.5 (E). Values above 10 or below 0.001 were plotted on the axis.

Finally, we analyzed Omicron immune-escape in the context of our knowledge of the neutralization potential of 163 out of the 313 monoclonal antibodies tested by BLI (Sokal et al., 2021b). As previously described for the Beta variants, loss of affinity against Omicron RBD appeared mostly restricted to potent neutralizers of the D614G SARS-CoV-2 (**Figure 2D**), highlighting the selective pressure imposed on SARS-CoV-2 by such antibodies. Moreover, antibodies that still displayed a potent neutralization potential against the Beta SARS-CoV-2 appeared selectively targeted for additional loss of affinity by Omicron-specific mutations not included in the Beta variant (**Figure 2E**).

## Discussion

MBCs display a diverse repertoire allowing for an adaptive response upon re-exposure to the pathogen, especially in the case of variants (Purtha et al., 2011; Weisel et al., 2016). We recently showed that the MBC pool elicited after mRNA vaccination in recovered and naive individuals displayed high-affinity neutralizing clones towards multiple VOCs, predicting the efficacy of the vaccine recall to cross-neutralize VOCs. We show here that the unique accumulation of mutations in key amino acid residues within the RBD of the Omicron variant results in a loss of affinity of 70% MBCs clones, greatly bypassing the Wuhan (WT) RBD-driven affinity-maturation process occurring in the germinal centers of infected and/or vaccinated individuals. Remarkably, it appears from our results that additional mutations selected in the Omicron variant might represent key RBD residues targeted by a large fraction of the potent anti-Beta SARS-CoV-2 neutralizing antibodies. As such, whereas high-affinity clones against Omicron were detectable in all donors, even in the relatively immature pool of naive vaccinated individuals, few of the potent anti-Beta SARS-CoV-2 neutralizing antibodies that we had previously described appeared unaffected in their recognition of the Omicron RBD (15 monoclonal antibodies out of a total of 164 tested for neutralization activity against Beta SARS-CoV-2, found in 5 out of the 9 individuals from which cells were tested). Given the limited number of fully tested antibodies per individuals (below 20 in 5 donors), it is difficult to give a precise range but one would expect that the available neutralizing MBC repertoire against the Omicron variant could fall closer to (or below) 10% of the overall anti-RBD MBC repertoire in most individuals. This low frequency of available protective MBC repertoire would represent a marked decrease compared to the 40 to 80% detected against the original SARS-CoV-2 in most donors (Sokal et al., 2021b). These results are consistent with a non-peer reviewed preprint analyzing sera collected from individuals that had received a mRNA recall vaccination mobilizing the MBC pool, either a third vaccine dose in naive individuals or a unique dose in previously infected individuals. In both cases, a more than 5-fold reduction in neutralization efficacy against the Omicron variant was measured as compared to the Wuhan (WT) and Delta strains. Nonetheless, given the extremely high serum antibody levels in these patients, all sera still displayed potent neutralizing activity against SARS-CoV-2 Omicron (Planas et al., 2021). Unlike plasma cells producing serum antibodies, MBCs are endowed with great proliferative potential and, at a rate of one division every 10-12 hours, could theoretically compensate for an up to 8-fold loss in protective clone frequency in less than 2 days, and may therefore be sufficient to avoid severe forms of COVID-19. Deeper analysis of the MBC repertoire and specific testing of the Omicron neutralization potential of memory B cell-derived antibodies from various individuals thus appear an important next step to better characterize the exact frequency of available protective MBCs against this latest SARS-CoV-2 variant.

A limitation to our conclusions relies in the fact that samples analyzed were collected early after the first vaccine boost in all individuals. The repertoire of COVID-19-naive vaccinated individuals has been shown to evolve up to 5 months after the boost (Cho et al., 2021; Goel et al., 2021a) and it remains to be assessed how this MBC repertoire will further evolve after an additional vaccine boost, with a third dose of the Wuhan (WT) spike mRNA vaccine currently being implemented in numerous countries. Additionally, our study was focused on the RBD domain of the SARS-CoV-2 spike protein, which represents the major target of neutralizing antibodies. However, neutralizing antibodies against other domains of the trimeric spike have been described, notably against the N-terminal domain (NTD) and these might be affected in a different way (Tong et al., 2021). A similar point could be made for memory T cell responses, which appear so far to be less affected by mutations selected by SARS-CoV-2 variants (Goel et al., 2021b). Answering these questions will provide us with crucial information regarding the available immune protection against Omicron or subsequent variants, and allow an informed decision as to whether vaccine specifically targeted against VOCs will need to be implemented in the near future.

## Acknowledgments

We thank Garnett Kelsoe for providing us with the human cell culture system, together with invaluable advices. We thank A. Boucharlat and the Chemogenomic and Biological screening core facility headed by F. Agou, as well as P. England and the Molecular Biophysics core facility at the Institut Pasteur, Paris, France for support during the course of this work. We also thank Sébastien Storck, Lucie Da Silva, Sandra Weller for their advices and support; we thank the physicians, Constance Guillaud, Raphael Lepeule, Frédéric Schlemmer, Elena Fois, Henri Guillet, Nicolas De Prost, Pascal Lim, whose patients were included in this study.

## RESOURCE AVAILABILITY

### Lead Contact

Further information and requests for resources and reagents should be directed to and will be fulfilled by the Lead Contact, Matthieu Mahévas.

### Materials Availability

No unique materials were generated for this study.

### Data and Code Availability

- Single-cell culture VDJ sequencing data are reported in Sokal et al.,2021b and included in this study as part of Table S1
- This paper does not report original code.
- Any additional information required to reanalyze the data reported in this paper is available from the lead contact upon request.

## EXPERIMENTAL MODEL AND SUBJECT DETAILS

### Study participants

In total, 43 patients with recovered COVID-19 (17 S-CoV and 26 M-CoV) and 25 naive patients were included in our previous studies and longitudinally followed-up after mRNA vaccination. SARS-CoV-2 infection was defined as confirmed reverse transcriptase polymerase chain reaction (RT-PCR) on nasal swab or clinical presentation associated with typical aspect on CT-scan and/or serological evidence. Naive patients were healthcare workers who had no history of COVID-19 and negative IgG anti-nucleocapsid (and/or Spike) (2018-A01610-55).

All vaccinated subjects received the BNT162b2 mRNA vaccine. SARS-CoV-2 recovered patients received only one dose, in line with French guidelines and naïve subject received two doses.

In this study, we included all the 9 patients (4 S-CoV, 4 M-CoV and 3 Naives patients) whose memory B cells were analyzed in Sokal et al. 2021b

Detailed information on the individuals included in this study, including gender and health status, can be found in Table S1.

Patients were recruited at the Henri Mondor University Hospital (AP-HP), between March and April 2021. MEMO-COV-2 study (NCT04402892) was approved by the ethical committee Ile-de-France VI (Number: 40-20 HPS), and was performed in accordance with the French law. Written informed consent was obtained from all participants.

## METHOD DETAILS

### Recombinant protein purification

#### Construct design

The SARS-CoV-2 Receptor Binding Domain (RBD) was cloned in pcDNA3.1(+) encompassing the Spike (S) residues 331-528, and it was flanked by an N-terminal IgK signal peptide and a C-terminal Thrombin cleavage site followed by Hisx8, Strep and Avi tags. The mutations present on the B.1.1.7 (Alpha, N501Y), B.1.351 (Beta, K417N, E484K, N501Y), P.1 (Gamma, K417T, E484K, N501Y), B.1.617.1 (Kappa, L452R, E484Q), and B.1.617.2 (L452R, T478K) variants as compared to the ancestral Wuhan (WT) strain were introduced by PCR mutagenesis using standard methods. B.1.1.529 (omicron variant) RBD plasmid was specifically designed with its mutations (N501Y, K417T/N, E484K/Q/A, T478K, G339D, S371L, S373P, S375F, N440K, G446S, S477N, Q493R, G496S, Q498R and Y505H).

#### Protein expression and purification

The plasmids coding for recombinant proteins were transiently transfected in Expi293F™ cells (Thermo Fischer) using FectoPRO^®^ DNA transfection reagent (Polyplus), according to the manufacturer’s instructions. The cells were incubated at 37 °C for 5 days and then the culture was centrifuged and the supernatant was concentrated. The proteins were purified from the supernatant by affinity chromatography using StrepTactin columns (IBA) (SARS-CoV-2 S) or His-Trap™ Excel columns (Cytiva) (SARS-CoV-2 RBD). A final step of size-exclusion chromatography (SEC) in PBS was also performed, using either a Superose6 10/300 column (Cytiva) for the SARS-CoV-2 S, or a Superdex200 10/300 (Cytiva) for the SARS-CoV-2 RBD.

### Single-cell culture

All culture supernatants from single-cell cultured memory B cells used in this study were generated and reported as part as a previous study (see Sokal et al., 2021b). Briefly, single B cells were sorted in 96-well plates containing MS40L^lo^ cells expressing CD40L (kind gift from G. Kelsoe, Luo et al., Blood 2009, Crickx et al., 2021). Cells were co-cultured at 37°C with 5% CO2 during 21 or 25 days in RPMI-1640 (Invitrogen) supplemented with 10% HyClone FBS (Thermo Scientific), 55 μM 2-mercaptoethanol, 10 mM HEPES, 1 mM sodium pyruvate, 100 units/mL penicillin, 100 μg/mL streptomycin, and MEM non-essential amino acids (all Invitrogen), with the addition of recombinant human BAFF (10 ng/ml), IL2 (50 ng/ml), IL4 (10 ng/ml), and IL21 (10 ng/ml; all Peprotech). Part of the supernatant was carefully removed at days 4, 8, 12, 15 and 18 and the same amount of fresh medium with cytokines was added to the cultures. After 21 days of single cell culture, supernatants were harvested and stored at −20°C. Cell pellets were placed on ice and gently washed with PBS (Gibco) before being resuspended in 50 μL of RLT buffer (Qiagen) supplemented with 10% 2-mercaptoethanol and subsequently stored at −80°C until further processing.

### ELISA

Total IgG from culture supernatants were detected by home-made ELISA. Briefly, 96 well ELISA plates (Thermo Fisher) were coated with goat anti-human Ig (10 μg/ml, Invitrogen) in sodium carbonate during 1h at 37°C. After plate blocking, cell culture supernatants were added for 1hr, then ELISA were developed using HRP-goat anti-human IgG (1 μg/ml, Immunotech) and TMB substrate (Eurobio). OD450 and OD620 were measured and Ab-reactivity was calculated after subtraction of blank wells.

### Single-cell IgH sequencing

Clones whose culture had proven successful (IgG concentration ≥ 1 μg/mL at day 21-25) were selected and extracted using the NucleoSpin96 RNA extraction kit (Macherey-Nagel) according to the manufacturer’s instruction. A reverse transcription step was then performed using the SuperScript IV enzyme (ThermoFisher) in a 14 μl final volume (42°C 10 min, 25°C 10 min, 50°C 60 min, 94°C 5 min) with 4 μl of RNA and random hexamers (Thermofisher scientific). A PCR was further performed based on the protocol established by Tiller et al (Tiller et al., 2008). Briefly, 3.5 μl of cDNA was used as template and amplified in a total volume of 40 μl with a mix of forward L-VH primers (**Table S1**) and reverse Cγ primer and using the HotStar® Taq DNA polymerase (Qiagen) and 50 cycles of PCR (94°C 30 s, 58°C 30 s, 72°C 60 s). PCR products were sequenced with the reverse primer CHG-D1 and read on ABI PRISM 3130XL genetic analyzer (Applied Biosystems). Sequence quality was verified with the CodonCode Aligner software (CodonCode Corporation).

### Computational analyses of VDJ sequences

Processed FASTA sequences from cultured single-cell VH sequencing were annotated using Igblast v1.16.0 against the human IMGT reference database. Clonal cluster assignment (DefineClones.py) and germline reconstruction (CreateGermlines.py) was performed using the Immcantation/Change-O toolkit (Gupta et al., 2015) on all heavy chain V sequences. Sequences that had the same V-gene, same J-gene, including ambiguous assignments, and same CDR3 length with maximal length-normalized nucleotide hamming distance of 0.15 were considered as potentially belonging to the same clonal group. Further clonal analyses on all productively rearranged sequences were implemented in R. Mutation frequencies in V genes were calculated using the calcObservedMutations() function from the Immcantation/SHazaM v1.0.2 R package. VH repartition was calculated using the countGenes() function from the Immcantation/alakazam v1.1.0 R package.

### 3D representation of known mutations to the RBD surface

Panel D in Figure 5 was prepared with The PyMOL Molecular Graphics System, Version 2.1 Schrödinger, LLC. The atomic model used for the RBD was extracted from the cryo-EM structure of the SARS-CoV-2 spike trimer (PDB:6XR8; Cai et al.,2020)

### Affinity measurement using biolayer interferometry (Octet)

Affinity measurement against the Wuhan (WT), B.1.1.7 (Alpha), B.1.351 (Beta), P.1 (Gamma), B.1.617.1 (Kappa) and B.1.617.2 (Delta) RBD were measured and reported as part of a previous study (see Sokal et al.,2021b). Affinity measurement against the B.1.1.529 Omicron RBD was performed in the same way, using new biosensors. As an internal quality control step to ensure the correct conservation of the supernatants a new measurement of the WT RBD affinity (WT2 in Table S1). All affinity measurement were done using biolayer interferometry assays on the Octet HTX instrument (ForteBio). This high-throughput kinetic screening of supernatants using single antigen concentration has recently been extensively tested and demonstrated excellent correlation with multiple antigen concentration measurements (Lad et al., 2015). Briefly, anti-Human Fc Capture (AHC) biosensors (18-5060) were immersed in supernatants from single-cell memory B cell culture (or control monoclonal antibody) at 25°C for 1800 seconds. Biosensors were equilibrated for 10 minutes in 10x PBS buffer with surfactant Tween 20 (Xantec B PBST10-500) diluted 1x in sterile water with 0.1% BSA added (PBS-BT) prior to measurement. Association was performed for 600 s in PBS-BT with WT or variant RBD at 100nM followed by dissociation for 600s in PBS-BT. Biosensor regeneration was performed by alternating 30s cycles of regeneration buffer (glycine HCl, 10 mM, pH 2.0) and 30s of PBS-BT for 3 cycles. Traces were reference sensor subtracted and curve fitting was performed using a local 1:1 binding model in the HT Data analysis software 11.1 (ForteBio). Sensors with response values (maximum RBD association) below 0.1nm were considered non-binding. For variant RBD non-binding mAbs, sensor-associated data (mAb loading and response) were manually checked to ensure that this was not the result of poor mAb loading. For binding clones, only those with full R^2^>0.8 were retained for KD reporting and initial prediction of key binding residues. mAbs were defined as affected against a given variant RBD if the ratio of calculated KD value against that RBD variant and the WT RBD was superior to two. Only monoclonal antibodies displaying correct loading (>0.4nm) and binding (>0.1nm) in our latest WT RBD affinity measurement (313/414) were retained in the analysis presented in this report.

Key binding residues predictions were extracted from our previous study (see Sokal et al.,2021b). Briefly, prediction were simply made based on mutations repartition in the different variants tested prior to the Omicron, for example mAbs affected only by B.1.351 and P.1 variants were predicted to bind to the K417 residue. Two exceptions to these simple rules were made: 1/ mAbs affected by the B.1.1.7, B.1.351 and P.1 variants were initially labeled as binding to the N501 residue but the ratios of KD values against the B.1.351 and P. 1 RBD variants and the B.1.1.7 RBD variants were further calculated. All mAbs with ratio superior to two for these two combinations were labeled as binding both the N501 and K417 residues, as previously described for RBS-A type of anti-RBD mAbs (Yuan et al., 2021); 2/ mAbs affected by the B.1.351, P.1, B.1.617.1 and B.1.617.2 variants, but not the B.1.1.7 variant, were labeled as binding both the E484 and L452 residues based on reported data in the literature for RBS-B/C antibodies (Yuan et al., 2021; Starr et al., 2021). Sensors with missing values were manually inspected to resolve binding residues attribution, leaving only four with an unresolved profile. As part of this new analysis, Omicron-specific monoclonal antibodies that had not been previously defined as binding one of the residue shared by previous variants analyzed were defined as binding Omicron-specific mutated residues (**Figure 2B**).

### Virus neutralization assay

Virus neutralization against the D614G and B.1.351 (Beta) SARS-CoV-2 were measured and reported as part of a previous study (see Sokal et al.,2021b). Briefly, virus neutralization was evaluated by a focus reduction neutralization test (FRNT). Vero E6 cells were seeded at 2×10^4^ cells/well in a 96-well plate 24h before the assay. Two-hundred focus-forming units (ffu) of each virus were pre-incubated with B-cell clone supernatants for 1hr at 37°C before infection of cells for 2hrs. The virus/antibody mix was then removed and foci were left to develop in presence of 1.5% methylcellulose for 2 days. Cells were fixed with 4% formaldehyde and foci were revealed using a rabbit anti-SARS-CoV-2 N antibody (gift of Nicolas Escriou) and anti-rabbit secondary HRP-conjugated secondary antibody. Foci were visualized by diaminobenzidine (DAB) staining and counted using an Immunospot S6 Analyser (Cellular Technology Limited CTL). B-cell culture media and supernatants from RBD negative clones were used as negative control.

Percentage of virus neutralization was calculated as (100 - ((#foci sample / #foci control)*100)). For culture supernatants, two IgG concentrations (80 nM and 16 nM) were tested for each sample and each virus. Potent neutralizers were defined as >80% neutralization at 16 nM, weak neutralizer as neutralization <80% at 16 nM but >80% at 80 nM. Others were defined as non-neutralizing.

### Quantification and Statistical Analysis

Statistical analyses were all performed using GraphPad Prism 9.0 (La Jolla, CA, USA).

### Additional resources

ClinicalTrials.gov Identifier: MEMO-CoV2, NCT04402892.

### Excel table titles

Table S1. Human subject informations and affinity measurement. Related to Figures 1 and 2

## References

Barnes, C.O., West, A.P., Huey-Tubman, K.E., Hoffmann, M.A.G., Sharaf, N.G., Hoffman, P.R., Koranda, N., Gristick, H.B., Gaebler, C., Muecksch, F., et al. (2020). Structures of Human Antibodies Bound to SARS-CoV-2 Spike Reveal Common Epitopes and Recurrent Features of Antibodies. Cell 182, 828–842.e16.

Cho, A., Muecksch, F., Schaefer-Babajew, D., Wang, Z., Finkin, S., Gaebler, C., Ramos, V., Cipolla, M., Mendoza, P., Agudelo, M., et al. (2021). Anti-SARS-CoV-2 receptor-binding domain antibody evolution after mRNA vaccination. Nature 600, 517–522.

Crickx, E., Chappert, P., Sokal, A., Weller, S., Azzaoui, I., Vandenberghe, A., Bonnard, G., Rossi, G., Fadeev, T., Storck, S., et al. (2021). Rituximab-resistant splenic memory B cells and newly engaged naive B cells fuel relapses in patients with immune thrombocytopenia. Sci. Transl. Med. 13, eabc3961.

Dugan, H.L., Stamper, C.T., Li, L., Changrob, S., Asby, N.W., Halfmann, P.J., Zheng, N.-Y., Huang, M., Shaw, D.G., Cobb, M.S., et al. (2021). Profiling B cell immunodominance after SARS-CoV-2 infection reveals antibody evolution to non-neutralizing viral targets. Immunity. 54, 1290–1303.e7.

Gaebler, C., Wang, Z., Lorenzi, J.C.C., Muecksch, F., Finkin, S., Tokuyama, M., Cho, A., Jankovic, M., Schaefer-Babajew, D., Oliveira, T.Y., et al. (2021). Evolution of antibody immunity to SARS-CoV-2. Nature 591, 639–644.

Goel, R.R., Apostolidis, S.A., Painter, M.M., Mathew, D., Pattekar, A., Kuthuru, O., Gouma, S., Hicks, P., Meng, W., Rosenfeld, A.M., et al. (2021a). Distinct antibody and memory B cell responses in SARS-CoV-2 naïve and recovered individuals following mRNA vaccination. Sci Immunol 6, eabi6950.

Goel, R.R., Painter, M.M., Apostolidis, S.A., Mathew, D., Meng, W., Rosenfeld, A.M., Lundgreen, K.A., Reynaldi, A., Khoury, D.S., Pattekar, A., et al. (2021b). mRNA vaccines induce durable immune memory to SARS-CoV-2 and variants of concern. Science eabm0829.

Gupta, N.T., Vander Heiden, J.A., Uduman, M., Gadala-Maria, D., Yaari, G., and Kleinstein, S.H. (2015). Change-O: a toolkit for analyzing large-scale B cell immunoglobulin repertoire sequencing data. Bioinformatics 31, 3356–3358.

Lad, L., Clancy, S., Kovalenko, M., Liu, C., Hui, T., Smith, V., and Pagratis, N. (2015). High-throughput kinetic screening of hybridomas to identify high-affinity antibodies using bio-layer interferometry. J. Biomol. Screen. 20, 498–507.

Luo, X.M., Maarschalk, E., O’Connell, R.M., Wang, P., Yang, L., and Baltimore, D. (2009). Engineering human hematopoietic stem/progenitor cells to produce a broadly neutralizing anti-HIV antibody after in vitro maturation to human B lymphocytes. Blood 113, 1422–1431.

Planas, D., Saunders, N., Maes, P., Guivel-Benhassine, F., Planchais, C., Buchrieser, J., Bolland, W.-H., Porrot, F., Staropoli, I., Lemoine, F., et al. (2021). Considerable escape of SARS-CoV-2 variant Omicron to antibody neutralization. BioRxiv 2021.12.14.472630.

Rodda, L.B., Netland, J., Shehata, L., Pruner, K.B., Morawski, P.A., Thouvenel, C.D., Takehara, K.K., Eggenberger, J., Hemann, E.A., Waterman, H.R., et al. (2021). Functional SARS-CoV-2-Specific Immune Memory Persists after Mild COVID-19. Cell 184, 169–183.e17.

Sokal, A., Chappert, P., Barba-Spaeth, G., Roeser, A., Fourati, S., Azzaoui, I., Vandenberghe, A., Fernandez, I., Meola, A., Bouvier-Alias, M., et al. (2021a). Maturation and persistence of the anti-SARS-CoV-2 memory B cell response. Cell 184, 1201–1213.e14.

Sokal, A., Barba-Spaeth, G., Fernández, I., Broketa, M., Azzaoui, I., Selle, A. de L., Vandenberghe, A., Fourati, S., Roeser, A., Meola, A., et al. (2021b). mRNA vaccination of naive and COVID-19-recovered individuals elicits potent memory B cells that recognize SARS-CoV-2 variants. Immunity 54, 2893–2907.e5.

Starr, T.N., Greaney, A.J., Dingens, A.S., and Bloom, J.D. (2021). Complete map of SARS-CoV-2 RBD mutations that escape the monoclonal antibody LY-CoV555 and its cocktail with LY-CoV016. Cell Rep Med 2, 100255.

Tong, P., Gautam, A., Windsor, I.W., Travers, M., Chen, Y., Garcia, N., Whiteman, N.B., McKay, L.G.A., Storm, N., Malsick, L.E., et al. (2021). Memory B cell repertoire for recognition of evolving SARS-CoV-2 spike. Cell 184, 4969–4980.e15.

Wang, Z., Schmidt, F., Weisblum, Y., Muecksch, F., Barnes, C.O., Finkin, S., Schaefer-Babajew, D., Cipolla, M., Gaebler, C., Lieberman, J.A., et al. (2021). mRNA vaccine-elicited antibodies to SARS-CoV-2 and circulating variants. Nature 592, 616–622.

Yuan, M., Huang, D., Lee, C.-C.D., Wu, N.C., Jackson, A.M., Zhu, X., Liu, H., Peng, L., Gils, M.J. van, Sanders, R.W., et al. (2021). Structural and functional ramifications of antigenic drift in recent SARS-CoV-2 variants. Science. 373, 818–823

Zahradník, J., Marciano, S., Shemesh, M., Zoler, E., Harari, D., Chiaravalli, J., Meyer, B., Rudich, Y., Li, C., Marton, I., et al. (2021). SARS-CoV-2 variant prediction and antiviral drug design are enabled by RBD in vitro evolution. Nat Microbiol 6, 1188–1198.

